# Novel Co-expression of Group I Metabotropic Glutamate Receptor During Mouse Cochlear Nucleus Development

**DOI:** 10.64898/2026.09.29.755315

**Authors:** Hitomi Sakano, Jason Wheeler, Justine Lam, Erik Vo, Robin Gibson, Jake Langevin, Xiaoyan Yu, Edwin W Rubel, Yuan Wang

## Abstract

Metabotropic glutamate receptors (mGluR) are G-protein coupled receptors that regulate neuroplasticity and have been shown to play a role in auditory processing. However, not much is known about the expression of mGluRs during cochlear nucleus development in the context of Fragile X syndrome (FXS). FXS is the most common heritable form of intellectual disability and autism and is associated with auditory processing deficits including hypersensitivity. Audiogenic seizure is a notable feature of the *Fmr1* knock-out (KO) mouse model for FXS, in which mGluR5 has been shown to be necessary. In this study, we investigated the expression of eight metabotropic glutamate receptors (mGluRs) in the cochlear nucleus at both young and adult ages and in wildtype and *Fmr1* KO mice. While it is established that mGluR1 is the predominantly expressed mGluR in adult cochlear nucleus, we report the following new finding that mGluR5 is the more abundant receptor during early development. The expression of Group II and III mGluRs (2,3,4,6,7,8) was relatively stable during development. We show that mGluR1 and mGluR5 are co-expressed across multiple neuron types. *Fmr1* genotype did not have statistically significant impact on the expression pattern of mGluRs in cochlear nucleus. Although hearing onset occurs during this mGluR5 to 1 transition period, mGluR expression patterns were not impacted by cochlear activity. We conclude that there is an mGluR5 is to mGluR1 transition that occurs within cochlear nucleus neurons during the critical period closure that is intrinsic to neuronal development.

## Introduction

Metabotropic glutamate receptors (mGluR) play an important role in modulating the auditory brainstem as well as many brain regions. In particular, mGluR5 has been shown to play an important role in the auditory processing deficits in Fragile X syndrome (FXS) which is the most common hereditary cause of autism. FXS is caused by a trinucleotide repeat expansion in the 5’ UTR of the *Fmr1* gene on the X-chromosome which leads to the methylation and silencing of the *Fmr1* gene and, as a result, loss of Fragile X messenger ribonucleoprotein 1 (FMRP) expression (Santoro et al., 2012). The *Fmr1* knock-out (KO) mouse has been widely used for mechanistic investigations of FXS neuropathology (Consortium, 1994). Auditory processing abnormalities have been described in individuals with FXS as well as in *Fmr1* KO rodents, including auditory hypersensitivity, altered pre-pulse inhibition of acoustic startle response, and increased susceptibility to audiogenic seizures to loud noise >120dB (Chen & Toth, 2001; Frankland et al., 2004; McCullagh et al., 2020; Nielsen et al., 2002; Rotschafer & Razak, 2014; Yun et al., 2006).

Signal pathways mediated by metabotropic glutamate receptors (mGluRs), particularly mGluR5, have long been proposed as key molecular players responsible for many forms of neuronal change induced by FMRP absence (Dolen & Bear, 2008; Huber et al., 2002; Thomazeau et al., 2021). Although earlier studies reported no or only marginal changes in mGluR5 level in the forebrain and cerebellum of rodents, more recent studies revealed altered mGluR5 availability in brains of FXS individuals and *Fmr1* KO mice using positron emission tomography (PET) imaging (Afshar et al., 2022; Giuffrida et al., 2005; Lohith et al., 2013; Mody et al., 2021). Consistent with this idea, adult mGluR5 KO rats display several FXS phonotypes including neuroinflammation and extracellular matrix changes (Dahl et al., 2022). Potential involvement of other mGluR receptors, however, remains unclear.

In the auditory system, multiple lines of evidence have linked abnormal mGluR5 signals to *Fmr1* KO-induced auditory deficits. Pharmacological mGluR5 inhibition normalized dendritic spine morphology and reduced the susceptibility of audiogenic seizures in *Fmr1* KO mice (Dolen et al., 2007; Gonzalez et al., 2019; Michalon et al., 2012; Yan et al., 2005). It is unknown, however, how mGluR expression is disrupted in the auditory system. Particularly, several auditory deficits, including enhanced susceptibility of audiogenic seizures and altered phase-locking, involve FMRP loss at the brainstem and midbrain levels (Gonzalez et al., 2019; Holley et al., 2022). Thus, in this study, we aimed to determine and compare the developmental expression of various metabotropic glutamate receptors in the cochlear nucleus of wildtype (WT) and F*mr1* KO mice.

The mGluR family consists of G-protein coupled receptors which are activated by glutamate. They are categorized into three functional groups that trigger distinct downstream signals. Group I (mGluR5 and mGluR1) receptors couple to the Gαq protein and activate phospholipase C pathways and intracellular calcium release. Group II (mGluR2 and mGluR3) and Group III (mGluR4, 6, 7 and 8) receptors couple to the Gαi/o protein and inhibit adenyl cyclase activity, leading to a decrease in intracellular cAMP levels (Niswender & Conn, 2010). It has been reported that mGluR1 is expressed in the adult mouse cochlear nucleus (Bilak & Morest, 1998; Lu, 2014; Petralia et al., 2000), while the expression pattern of other mGluR receptors and their developmental trajectories have not been investigated. Here, we present novel mGluR expression data in the cochlear nucleus of the WT and *Fmr1* KO mice during postnatal development, by combining RNA (RNA sequencing and RT-PCR) and protein (immunohistochemistry and western blot) analyses. We identified a transition from predominantly mGluR5 to mGluR1 expression during the second postnatal week at the time of hearing onset and closure of the critical period. However, this transition did not require cochlear input. We also did not detect a significant difference in mGluR expression levels between WT and *Fmr1* KO mice.

## Material and Methods

### Animals and tissue collection for RNA sequencing

All animal procedures were approved by the Institutional Animal Care and Use Committees at University of Texas Southwestern Medical Center, University of Rochester Medical Center, University of Washington, and Florida State University (FSU) and were conducted in accordance with the National Institutes of Health policies on animal use. All mouse breeders were obtained from the Jackson Laboratory (Bar Harbor, MA, USA) and housed in institutional animal facilities for breeding. The day of birth was defined as postnatal day 0 (P0).

For comparing mGluR expression between WT and *Fmr1* KO, WT (C57Bl/6J, RRID: IMSR_JAX:000664) and *Fmr1* KO mice (B6.129P2-*Fmr1^tm1Cgr^*/J, RRID: IMSR_JAX:003025) were maintained with intermittent backcrossing to C57Bl/6J. The cochlear nucleus (including both the dorsal and ventral portions) was dissected freshly at the age of P8 and two months (P60). For each genotype at each age, three biological replicates were collected, with each sample containing four cochlear nuclei from two animals.

For afferent deprivation in DTR mice, WT and DTR mice were injected with DT (5 ng/g body weight) or 0.9% saline at age P4. Cochlear nuclei were collected at P7 and P21 for RNA sequencing analyses. For double-transgenic mice carrying DTR and *Fmr1* KO, heterozygote male *Pou4f3^DTR/^*^+^ mice (B6.Cg-*Pou4f3^TM1.1(HBEGF)Jsto^*/RubelJ; RRID: IMSR_JAX:028673) were crossed with either WT or *Fmr1* KO female mice. The resultant *Fmr1* KO:*Pou4f3^DTR/^*+ male pups from *Fmr1* KO dams were then crossed with *Fmr1* KO females. This breeding paradigm produced four offspring genotypes: *Pou4f3^DTR/^*^+^ (DTR), *Pou4f3^DT+/^*^+^ (WT), *Fmr1* KO:*Pou4f3^DTR/^*^+^ (*Fmr1* KO:DTR), and *Fmr1* KO: *Pou4f3^DTR/^*^+^ (*Fmr1* KO) (Yu & Wang, 2023). DTR and *Fmr1* KO:DTR mice were used for RNA-Seq analysis. Mice received a single injection of DT (5 ng/g body weight) or 0.9% saline intramuscularly at P5. DT induced hearing loss in DTR and *Fmr1* KO:DTR mice was previously confirmed by elevated thresholds of auditory brainstem responses (Yu & Wang, 2023). The cochlear nucleus was collected at P11 for RNA sequencing analyses. Briefly, mouse brains were rapidly extracted and placed in ice-cold phosphate-buffered saline (PBS). Coronal slices of tissue (∼ 2 mm thick) containing the entire cochlear nucleus were isolated, from which the cochlear nucleus from both sides were dissected and snap frozen in dry ice and stored at-80°C. Cochlear nuclei from 1-3 mice were pooled as one biological sample. Five biological replicates for each genotype and each treatment (DT vs. saline injection) were collected and used for RNA sequencing analyses.

### RNA sequencing

RNA was extracted by homogenization in Tris-buffered saline (TBS; 10mM TrisHCl, 150mM NaCl, 10mM EDTA, 0.5% Triton™ X-100) followed by purification using RNeasy Mini kits (QIAGEN; Venlo, Netherlands) and DNase I digestion following manufacturer’s instructions. RNA-seq libraries were constructed from 200 ng of RNA using the TruSeq Stranded mRNA Library Prep Kit (Illumina; San Diego, CA) following manufacturer’s instructions. The quantity and quality of the libraries were measured using a Qubit 4 Fluorometer (Thermo Fisher Scientific; Waltham, MA) and a 5300 Fragment Analyzer System (Agilent; Santa Clara, CA), respectively. The sequencing was performed on an Illumina NextSeq550 at the University of Rochester and on Illumina HiSeq 2000 at the University of Washington. Raw reads generated from the Illumina basecalls were demultiplexed using the bcl2fastq (version 2.19.1). Quality filtering and adapter removal were performed using the FastP (version 0.20.0) with parameters “--length_required 35 --cut_front_window_size 1 --cut_front_mean_quality 13 -- cut_front --cut_tail_window_size 1 --cut_tail_mean_quality 13 --cut_tail-w 8-y-r-j“. Cleaned reads were aligned to the Mus musculus reference genome (GRCm38.p6 + Gencode-M22 Annotation) using STAR_2.7.0f with parameters “ --twopassMode Basic --runMode alignReads --genomeDir ${GENOME} --readFilesIn ${SAMPLE} --outSAMtype BAM Unsorted -- outSAMstrandField intronMotif --outFilterIntronMotifs RemoveNoncanonical”. For RNA-seq analyses, gene-level read quantification was derived using the subread-1.6.4 package (featureCounts) with a GTF annotation file (Gencode M22) and parameters “-s 2-t exon-g gene_name”. For comparative transcript expression analyses, transcript-level reads were quantified with Salmon-0.13.1. Data normalization and differential expression analysis between WT and *Fmr1-*KO groups was performed using DESeq2-1.22.1 using median of ratios normalization, an adjusted P-value threshold of 0.05 within R version 3.5.1 (https://www.R-project.org/). Pheatmap (version 1.0.12; https://CRAN.R-project.org/package=pheatmap) was used to produce the heatmap of differentially expressed genes. The RNA-seq raw data is available through GEO (https://www.ncbi.nlm.nih.gov; accession numbers GSE236056, GSE346517, GSE346949 and GSE347368). Genes with log2 fold >0.1 and base count mean >100 were selected for gene ontology analysis with DAVID program (https://david.ncifcrf.gov/) (Huang da et al., 2009) and Syngo (https://www.syngoportal.org/).

Existing cochlear nucleus patch-seq raw data was obtained from GEO (GSE273145) (Jing et al., 2025) and processed using Seurat package in R. Gene expression matrices were normalized with NormalizeData with LogNormalize method and scale factor of 10000. Highly variable genes were identified with FindVriableFeatures with vst method and 2000 features. Data were scaled with ScaleData, followed by principle component analysis (RunPCA). Cell clusters were identified using shared nearest-neighbor graph with FindNeighbors on the first 10 principal components and FindClusters at a resolution of 0.5. RunUMAP was used to perform Uniform Manifold Approximation and Projection (UMAP) and DimPlot to visualize cell clusters. Expression of selected genes were also visualized violin plots using VlnPlot. Genetic markers of inhibitory and excitatory neurons, as well as neuron type-specific markers reported by Jing et al. were used to confirm appropriate clustering.

### RT-qPCR

RNAs were extracted from the dissected cochlear nuclei. Tissues were lysed in 6M urea. A portion of this lysis was extracted with 500ul TRIzol® (Invitrogen) with 100ul of choloroform added, vortexed, and set aside for 2 minutes. Samples were then centrifuged for 5 minutes at 15,000 x g at 4°C. The aqueous phase was precipitated with equal volume isopropanol and 5 µg glycogen carrier. Pellets were washed three times with 75% ethanol, centrifuged at 15,000 x g for 5 minutes, air dried for 5 minutes, and suspended in ultrapure RNase free water. DNase treatment was performed using RQ1 DNase per manufacturer instructions (Promega). RNA was re-extracted with TRIzol® as above and re-suspended in RNase-free water. RNA (250 ng) was reverse transcribed using qScript cDNA SuperMix (Quantabio), per manufacturer’s instructions, and diluted to 200 µl. A portion (2 µl) was then used in 10 µl quantitative (q)PCR reactions that employed SYBR™ Select Master Mix (Thermo Fisher Scientific). PCR was undertaken using a QuantStudio™ Real-Time PCR System (Thermo Fischer Scientific) using serial dilution control using WT samples. Quantitation was performed using the standard curve method and Pfaffl calculation. The following primer sets were utilized:

*mGluR5*: TGGGTCTAGCAGGGGAATTT; TGTAATCCCACCGACAGCTT

*mGluR1*: GACACCTCTTGGAAAACCCC; GCGTTGATGACAAATCCCAT

*ß-actin*: CACTGTCGAGTCGCGTCC; CCTTCTGACCCATTCCCACC

Cycling conditions: 94°C for 2 minutes, followed by 40 cycles at 94°C for 15 seconds and 60°C for 1 minute.

### Western blotting

Dissected cochlear nuclei were homogenized with a pestle in 6M urea, sonicated for 10 seconds, and then centrifuged at 14,000 x g for 10 minutes to remove cellular debris. Protein was quantitated using a Bradford Assay (Bio-Rad). Protein lysate was incubated using 1:5 volume of 5x Laemli buffer (250mM Tris-HCl pH 6.8, 30% glycerol, 10% sodium dodecyl sulfate, 0.02% bromophenol blue, 5% ß-mercaptoethanol) at 60°C for 30 minutes to prevent aggregation of mGluR protein. SDS-PAGE and western blotting were performed as described previously (Litovchick, 2020) using 10 µg of protein per lane. Blots were blocked in 5% milk in TBS-Tw (Tris-HCl 50mM pH 7.5, 150mM NaCl, 0.05% Tween-20). Blots were incubated with primary antibodies diluted in TBS-Tw overnight at 4°C, rinsed with TBS-Tw, incubated with horse radish peroxidase-conjugated secondary antibodies in TBS-Tw for one hour at room temperature and rinsed for 30 minutes in TBS-Tw. See **Table 1** for the primary antibodies used. Blots were developed using the Pierce™ SuperSignal™ West Pico Western Blot Kit (Thermo Scientific) and imaged using a ChemiDoc™ Touch Imaging System (Bio-Rad).

**Table 1.** Antibodies used in this study. IHC, immunohistochemistry. W, western blotting. aa, amino acid. AR, antigen retrieval. MW, molecular weight (on western blotting).

| Detects | Host, Dilution | MW (kDa) | Catalogue | RRID |
| --- | --- | --- | --- | --- |
| FMRP human aa550-C-term | Rabbit, polyclonal<br>1:1000 (W), 1:500 (IHC) | ~75<br>triplet | Abcam,<br>ab17722 | AB_2278530 |
| mGluR1 | Rabbit, monoclonal<br>1:500 (W) | ~150 | Cell Signaling<br>12551 | AB_2797953 |
| Rat mGluR1α | Mouse, monoclonal<br>1:150 (IHC AR) | | BD Biosciences<br>556389 | AB_396404 |
| mGluR5 | Rabbit, polyclonal<br>1:150 (IHC AR), 1:2000 (W) | ~150 | Abcam<br>ab76316 | AB_1523944 |
| Rabbit IgG | Goat, HRP-conjugated<br>polyclonal<br>1:30,000 (W) |  | Abcam<br>ab6721 | AB_955447 |

### Immunohistochemistry (IHC)

Mice were anesthetized and perfused with phosphate-buffered saline (PBS) (137mM NaCl, 2.7mM KCl, 10mM Na_2_HPO and 1.8 mM KH_2_PO4) followed by preservation in fixative (4% paraformaldehyde in PBS) for 2 hours, cryoprotection in 30% sucrose in PBS for 72 hours, and freezing in O.C.T. Medium (Sakura). Cryosections (14 µm) of the cochlear nucleus were collected using a cryostat (Leica CM1860 UV) and mounted on SuperFrost glass slides (Fisher Scientific). Slides were rehydrated in Tris-buffered saline (TBS) (100mM Tris HCl, 150mM NaCl, pH7.5). Slides were then incubated with TBS with 0.1% Triton™ X-100 and TNB Blocking Buffer (Perkin Elmer) for 30 minutes. See **Table 1** for the primary antibodies used. Antibody incubations were performed in TNB Blocking Buffer overnight at 4°C. Rabbit primary antibodies were detected with Alexa-488-conjugated goat secondary antibody at a 1:500 dilution (Invitrogen #A11034; RRID:AB_2576217). Mouse primary antibodies were detected using the M.O.M.® kit (Vector Labs) and Cy3-conjugated streptavidin (Jackson Immunoresearch, 1:1000). Following washing in TBS, slides were mounted using Fluoromount-G™ (Invitrogen) and imaged with a 20x objective using a Zeiss Palm MicroBeam microscope. Images were analyzed with the Image J software (version 2.14.0; NIH) (Schneider et al., 2012). For each WT and *Fmr1* KO case, four to six sections containing the main body of the VCN were analyzed. VCN from each section was imaged at an x-y resolution of 0.65 µm. Pixels above background intensity (measured at the adjacent spinal tract with low signal for mGluR) for each antibody (mGluR1 or mGluR5) were counted and the percentage of positive pixels was calculated.

### Antibody Characterization

The antibodies used for western blotting and IHC are listed in **Table 1**. The antibody specificities were confirmed by the bands at the expected molecular weights by western blotting. Antibodies against mGluR5 and FMRP were also confirmed with negative bands using tissues from mGluR5 KO (data not shown here) and *Fmr1* KO mice. Furthermore, the pattern of mGluR5 and mGluR1 immunostaining was confirmed with the developmental changes in staining levels that correlate with corresponding mRNA levels by *in situ* hybridization and RT-qPCR.

### RNAscope *in situ* hybridization

Brain cryosections were prepared from mice as described above for IHC with the exception that brains were postfixed for 24 hours. RNAscope was performed with RNAscope™ Multiplex Fluorescent Reagent Kit v2 according to the manufacturer protocol (Advanced Cell Diagnostics #323100). Probes for mouse mGluR5 (#423631) and mGluR1 (#449781), along with positive (#320881) and negative (#320871) control probes, were utilized.

### Statistical analyses

Statistical analyses were conducted using GraphPad Prism software (GraphPad, San Diego, CA, USA). One-way ANOVA followed by Tukey’s multiple comparison was used to compare normalized western band sizes and immunostaining intensities across ages in WT mice. Two-tailed, unpaired Student’s *t-*test was used to compare RT-PCR measures between ages in WT mice. Parametric test with Welch’s correction and non-parametric Mann-Whitney tests were used for data that pass and fail Shapiro–Wilk normality test, respectively. Two-way ANOVA followed by Šídák multiple comparisons were used to determine the effects and interaction between *Fmr1* genotype and age or afferent condition on mGluR1 and mGluR5 expression and protein level.

## Results

### A developmental switch in mGluR5-to-mGluR1 gene expression in cochlear nucleus

To compare gene expression levels of mGluRs, specifically mGluR1-8, we performed RNA sequencing (RNA-seq) on mRNAs isolated from the cochlear nucleus, containing both the dorsal and ventral portions (DCN and VCN), of WT and *Fmr1* KO mice at postnatal (P) day 8 (P8) and P60. Normalized base counts were analyzed for the most abundant transcript for each gene, averaged across biological replicates of the same age, and plotted as mean with standard deviations for each receptor (**Fig. 1A-B**). Consistent with previous reports (Bilak & Morest, 1998; Petralia et al., 2000; Tang & Lu, 2018), the most abundant mGluR in the WT adult (P60) cochlear nucleus was *mGluR1* (**Fig. 1A**, gray bars). The normalized base count of *mGluR1* was 3-5 folds higher than *mGluR3*, *4*, or 5, and more than 12 folds higher than *mGluR2*, *7*, or *8*. In the developing WT cochlear nucleus at P8, Group I receptors (*mGluR1* and *mGluR5*) collectively were also the most abundant mGluRs (**Fig. 1B**). However, *mGluR5* exhibited the highest expression level across the eight mGluRs, with normalized base counts being 2.5 folds higher than *mGluR1*. Relative abundance levels of Group II and III receptors were consistently low, showing similar patterns between P8 and P60. At both ages, progressive reductions were observed for *mGluR4*, *mGluR3*, *mGluR7*, *mGluR8, and mGluR2*, while *mGluR6* remained undetected.

**Figure 1.**
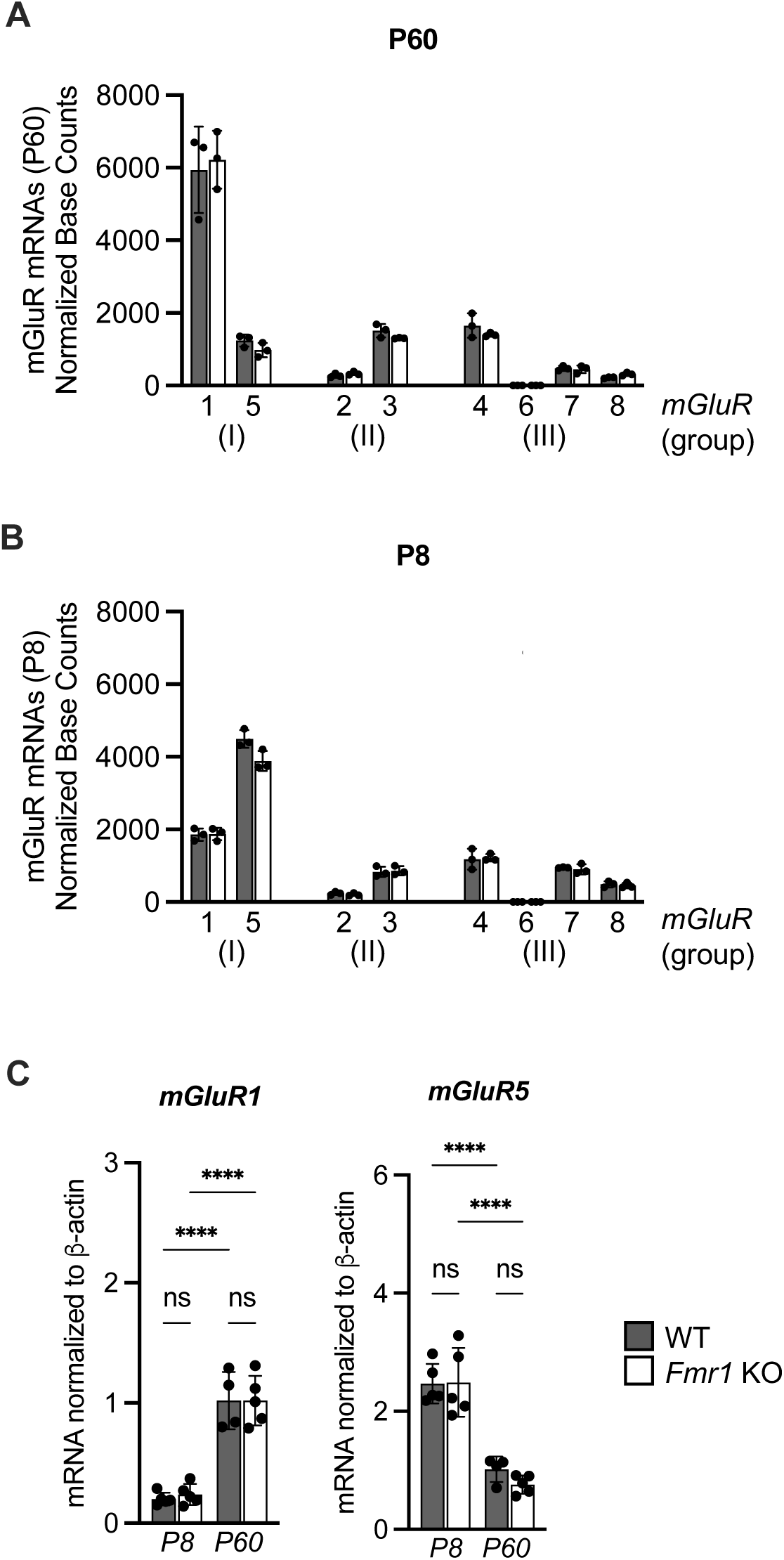
Group I *mGluR* expression in the cochlear nucleus changes with development. **(A-B)** For each *mGluR* (1-8) belonging to one of three group (I-III), the abundance of mGluR mRNA using normalized base mean count for the most abundant protein-coding isoform from the RNA-seq data are plotted for WT (gray bar) and *Fmr1-*KO (white bar) mice (n = 3, each) at postnatal day 60 (P60) and 8 (P8). At P60, *mGluR1* is most abundant while at P8, *mGluR5* is most abundant. (There is no significant genotype effect by Holm-Šídák multiple comparisons test, n = 3). **(C)** RT-qPCR confirmation of *mGluR1* and *mGluR5* levels at P8 and P60 are shown for both WT and *Fmr1-*KO mice. There was no difference between genotypes but there was significant difference between ages (****p < 0.0001, Two-way Anova and Fisher’s LSD test, n = 5 except for WT P60 where n = 4).

To validate age-dependent patterns of *mGluR1* and *mGluR5* expression, RT-qPCR was performed on samples collected from the cochlear nucleus (n = 5 biological replicates except for WT at P60 where n = 4) (**Fig. 1C**). *mGluR1* expression was over five-fold higher at P60 than at P8 (1.02 vs 0.20, *p* < 0.0001, two-way ANOVA, Fisher’s LSD), while *mGluR5* expression was over two-fold higher at P8 than P60 (2.47 vs 1.02, *p* < 0.0001, two-way ANOVA, Fisher’s LSD). These findings are consistent with the RNA-seq data.

There was no effect of *Fmr1* on mRNA expression of mGluR genes, comparing base counts between age-matched WT and *Fmr1* KO mice, at P8 or P60 for any of the mGluRs (Holm-Šídák multiple comparisons test) (**Fig. 1A-B**; white bars). The results are validated by RT-qPCR analysis for *mGluR1* (Two-way ANOVA; F_(1, 15)_ = 0.0599, *p* = 0.81) and *mGluR5* (F_(1, 15)_ = 0.4875, *p* = 0.4957) (**Fig. 1C**; white bars) where we see an effect of age but not with genotype.

### mGluR5-to-mGluR1 protein expression mirrors the mRNA expression and there is no statistically significant effect of *Fmr1* KO

We examined the developmental profiles of mGluR1 and mGluR5 at the protein level using western blotting on cochlear nucleus tissue (VCN and DCN combined) collected at P8, P15, P22, P29, and P90 from WT and *Fmr1* KO males. The developmental pattern of protein expression of mGluR5 and mGluR1 mirrored mRNA expression patterns (**Fig. 2A)**. In WT, average western blot band intensities for mGluR5 (normalized to β-actin) level was high at P8 and declined through P15 and remained low past P22 (**Fig. 2B**, dotted line), while mGluR1 level showed a steady increase with age (**Fig. 2B**, solid line) (n = 4). A similar pattern was observed in *Fmr1* KO mice (**Fig. 2C**) (n = 4). The same data are plotted to compare mGluR between WT and *Fmr1* KO. For mGluR5, there was an effect of age (F_(4,30)_ = 42.75, *p* < 0.0001) but not genotype (F_(1,30)_ = 2.073, *p* = 0.16) (**Fig. 2D**) by Two-way ANOVA. Posthoc Šídák’s multiple comparisons test shows statistically significant differences between P8 and each of the other ages (*p <* 0.0001). For mGluR1, there was main effect of age (F_(4,30)_ = 2.8, *p* < 0.05) but not genotype (F_(1,30)_ = 0.1, *p* = 0.68) (**Fig. 2E**). Posthoc analysis did not show significant differences between ages. We also performed western blotting analysis of Group1 mGluR expression normalized to total protein between WT and *Fmr1* KO mice across development for females in the C57Bl/6J strain, and males and females in the FVB strain with similar results (**Fig. 3**, n = 3-5 each).

**Figure 2.**
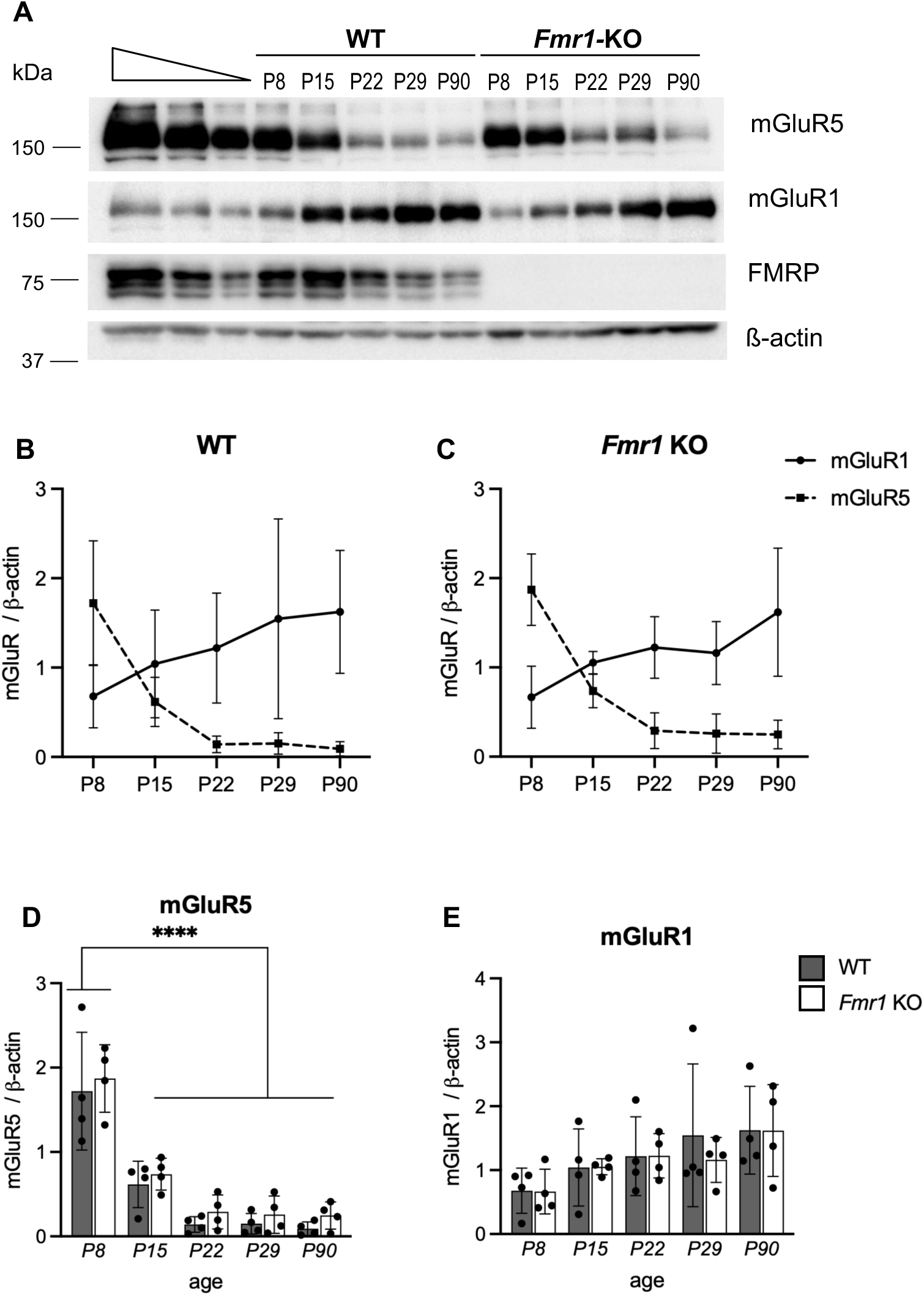
Western blotting shows an inverse pattern of mGluR5 and mGluR1 in cochlear nucleus across development. (A) Ventral and dorsal cochlear nuclei from WT and *Fmr1* KO male mice in the C57Bl6/6J strain of ages P8-P90 were analyzed for mGluR5 and mGluR1, with ß-actin as loading control. (B,C) Quantitation for protein band intensity normalized to ß-actin shows mGluR5 (dotted line) is highly expressed in both WT and KO mice during early development and decreases with age. There is an inverse pattern with mGluR1 (solid line). (D,E) There is a main effect of age for mGluR5 and for mGluR1 by Two-way ANOVA. Posthoc Šídák’s multiple comparisons test show significant difference of mGluR5 at P8 compared to other ages. (***p < 0.0001, n = 4 each).

**Figure 3.**
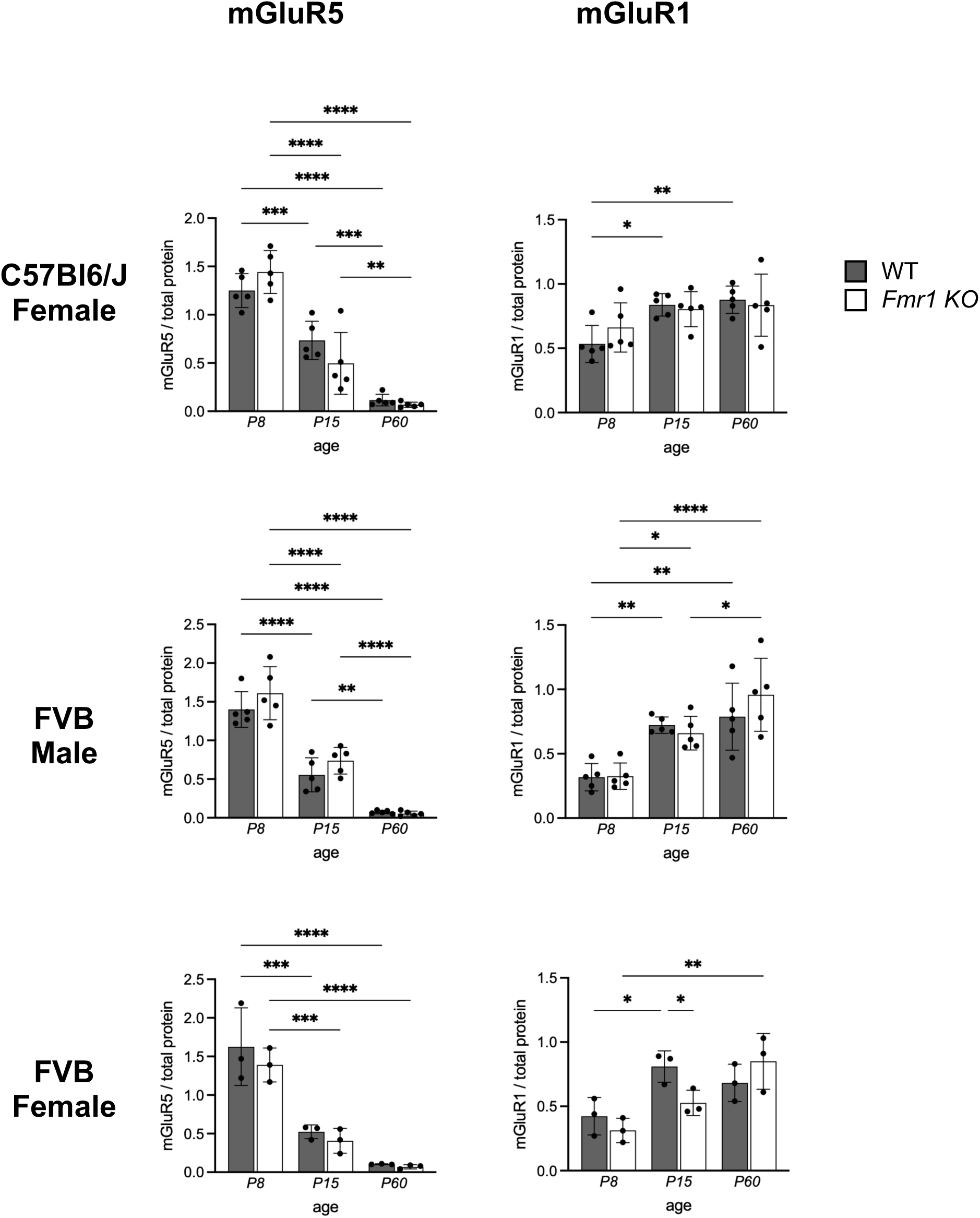
Western blotting for mGluR5 and mGluR1 of cochlear nuclei from females and additional strains across age. Quantitation of western blotting for mGluR5 (left column) and mGluR1 (right column) normalized to total protein stain, plotted by age for each WT (white bar) and *Fmr1* KO (gray bar) mice. For each graph, there is a main effect of age for mGluR5 and for mGluR1 by Two-way ANOVA. Posthoc Šídák’s multiple comparisons test show significant differences. (***p < 0.001, ****p < 0.0001, n = 5 each, n = 3 for FVB females).

Together, these findings illustrate a developmental transition in the cochlear nucleus, characterized by predominant mGluR5 expression during early postnatal stages followed by mGluR1 dominance in adulthood, a process referred to as the mGluR5-mGluR1 transition.

### Co-expression of mGluR5 and mGluR1

To examine the regional localization of mGluR1 and mGluR5 protein in the cochlear nucleus, we performed immunohistochemistry in anteroventral cochlear nucleus (AVCN) across ages at P8, P15, and P60 **(Fig. 4A**, outlined with dotted white line). There is a significant effect of age (F_(2, 12)_ = 11.06, *p* < 0.01, One-way ANOVA, posthoc Tukey’s test) between P8 and P60 on the ratio of the overall immunostaining intensity of mGluR5 to mGluR1 **(Fig. 4B).** As a second analysis, the percentages of the pixels (1 pixel = 0.65 x 0.65 μm) that were immunoreactive to mGluR1 only (red), mGluR5 only (green), both (yellow), or neither (black, i.e. below background threshold) were calculated **(Fig. 4C)**. Over half of the pixels within the AVCN showed overlapped staining (yellow) at each age (53% at P8, 58% at P15, and 59% at P60). The percentage of mGluR5-only pixels decreased with age (21% at P8, 17% at P15, and 12% at P60), while the percentage of mGluR1-only pixels increased with age (8% at P8, 11% at P15, 15% at P60). The same analyses were performed for PVCN and DCN (**Fig. 5A)**. As in AVCN, the majority of pixels were positive for both mGluR5 and mGluR1 (yellow), and there was a trend of decreasing mGluR5 to mGluR1 ratios with age, although it did not reach statistical significance (**Fig. 5B)**.

**Figure 4.**
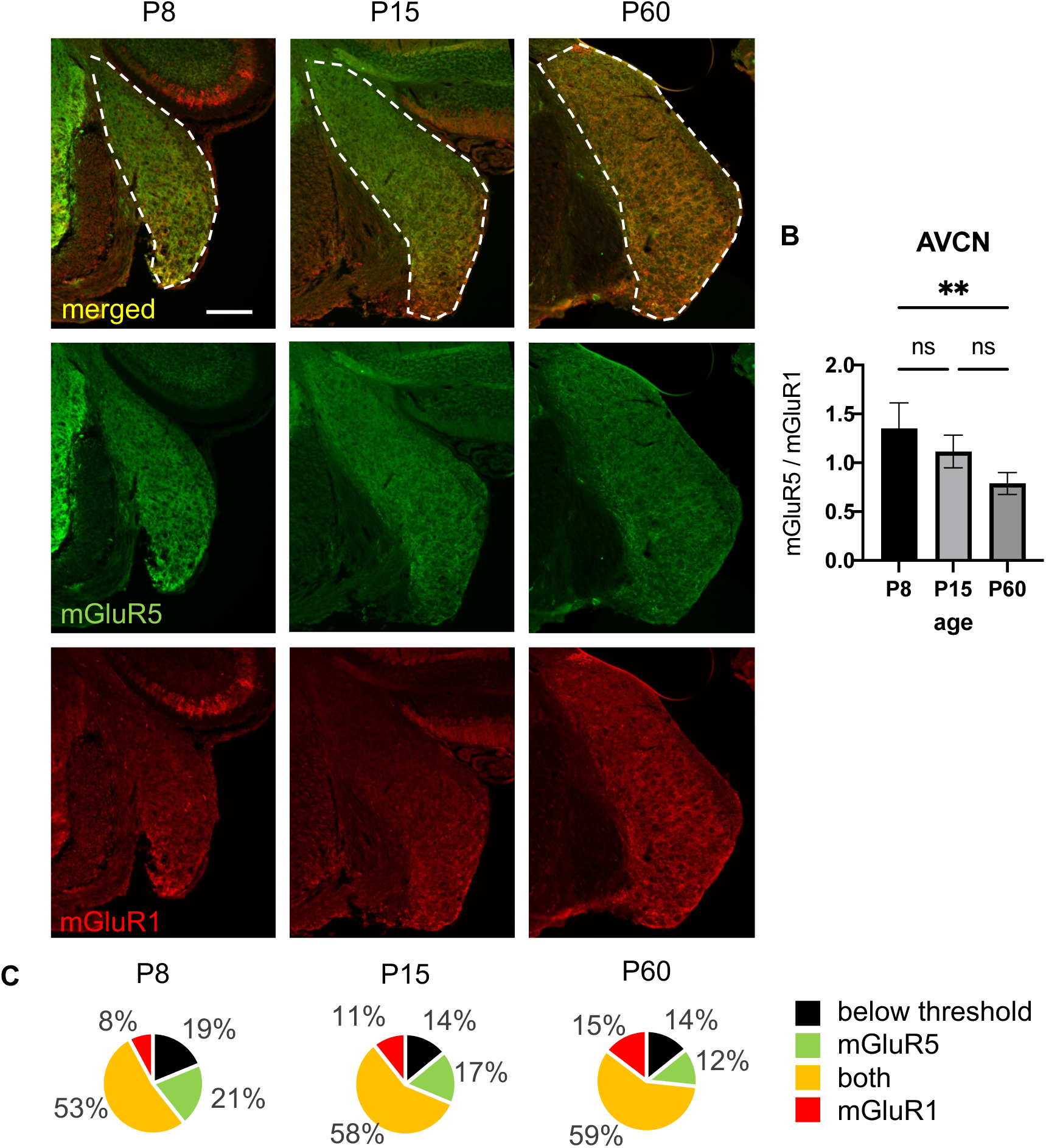
The distribution of mGluR5 and mGluR1 expression in the AVCN across age. (A) Shown are expression pattern of mGluR5 (green) and mGluR1 (red) at P8, P15 and P60 in the AVCN (outlined in dotted white line). The distribution of staining appears similar across ages. (B) The percentage of image pixels within the area of the AVCN for signal above background (obtained at the fibers of the adjacent trapezoid body) thresholds for mGluR5 (green), mGluR1 (red), both (yellow) or neither (black). The majority of pixels are co-labeled. Although not statistically significant, there is a slight trend of decreasing proportion of mGluR5 only pixels and increasing proportion of mGluR1 pixels with age. (C) The average pixel intensity ratio of mGluR5 to mGluR1 over the entire AVCN declines with age (n = 3 mice). **p < 0.01, one-way ANOVA with posthoc Tukey’s test, 1 pixel = 0.65 x 0.65µm, scale bar = 100µm.

**Figure 5.**
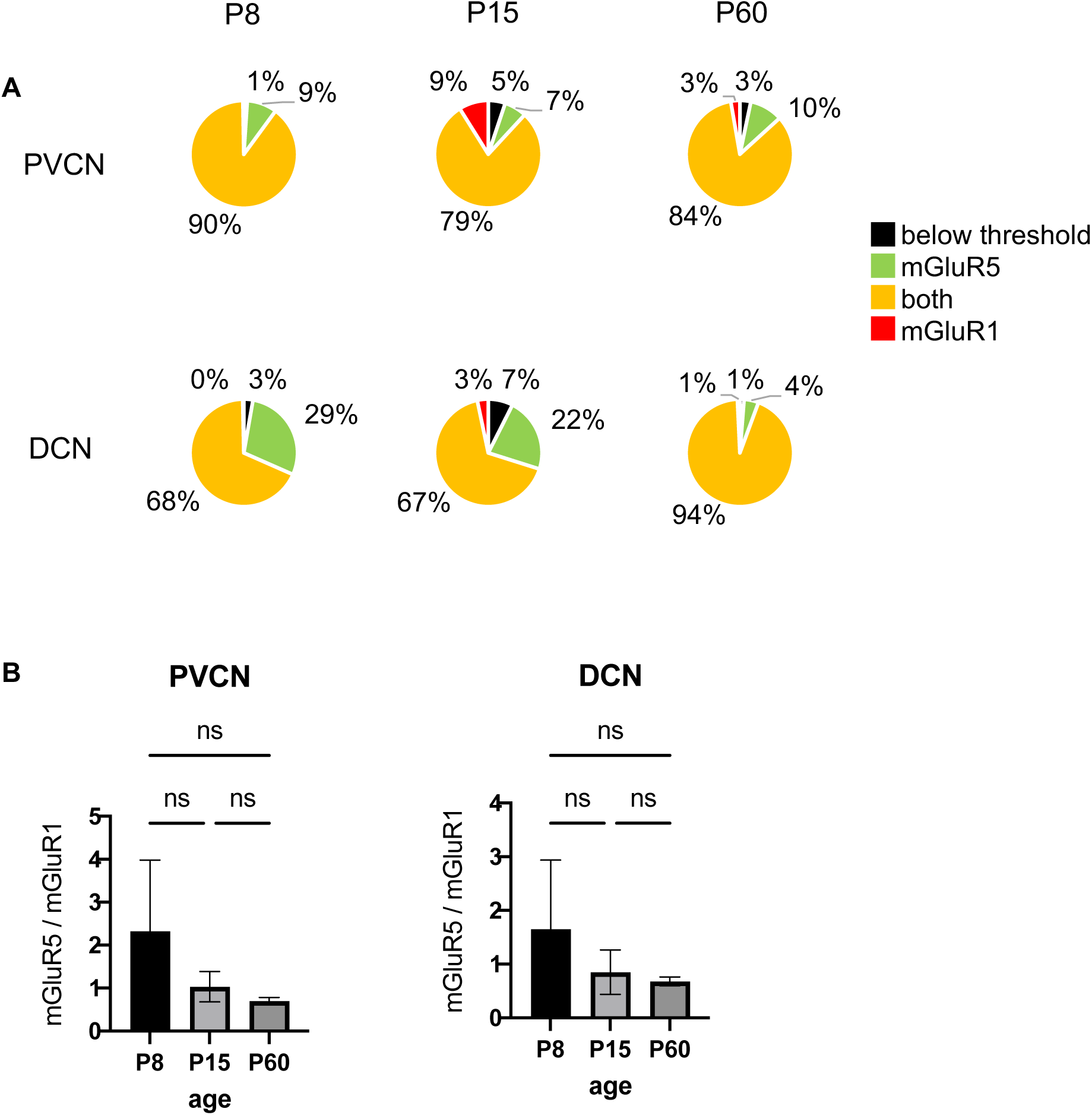
The distribution of mGluR5 and mGluR1 expression in the PVCN and DCN across age. (A) The percentage of image pixels within the area of the PVCN or DCN for signal above background thresholds for mGluR5 (green), mGluR1 (red), both (yellow) or neither (black). The majority of pixels are co-labeled. (B) The intensity of mGluR5 compared to mGluR1 within a cell trended down with age, although it did not reach statistical significance. (n = 3 mice).

The cells expressing either mGluR5 or mGluR1 in AVCN were counted for each age (P8, P15 and P60). At each age, the majority of cells (>75%) co-expressed both receptors (**Fig.6A**). The ratio of cytoplasmic mGluR5 to mGluR1 intensities decreased with age (*p* < 0.05, One-way ANOVA with posthoc Tukey’s test) (**Fig. 6B**). An example of confocal imaging of cells co-expressing both mGluR5 and mGluR1 at age P15 is shown (**Fig. 6C**).

**Figure 6.**
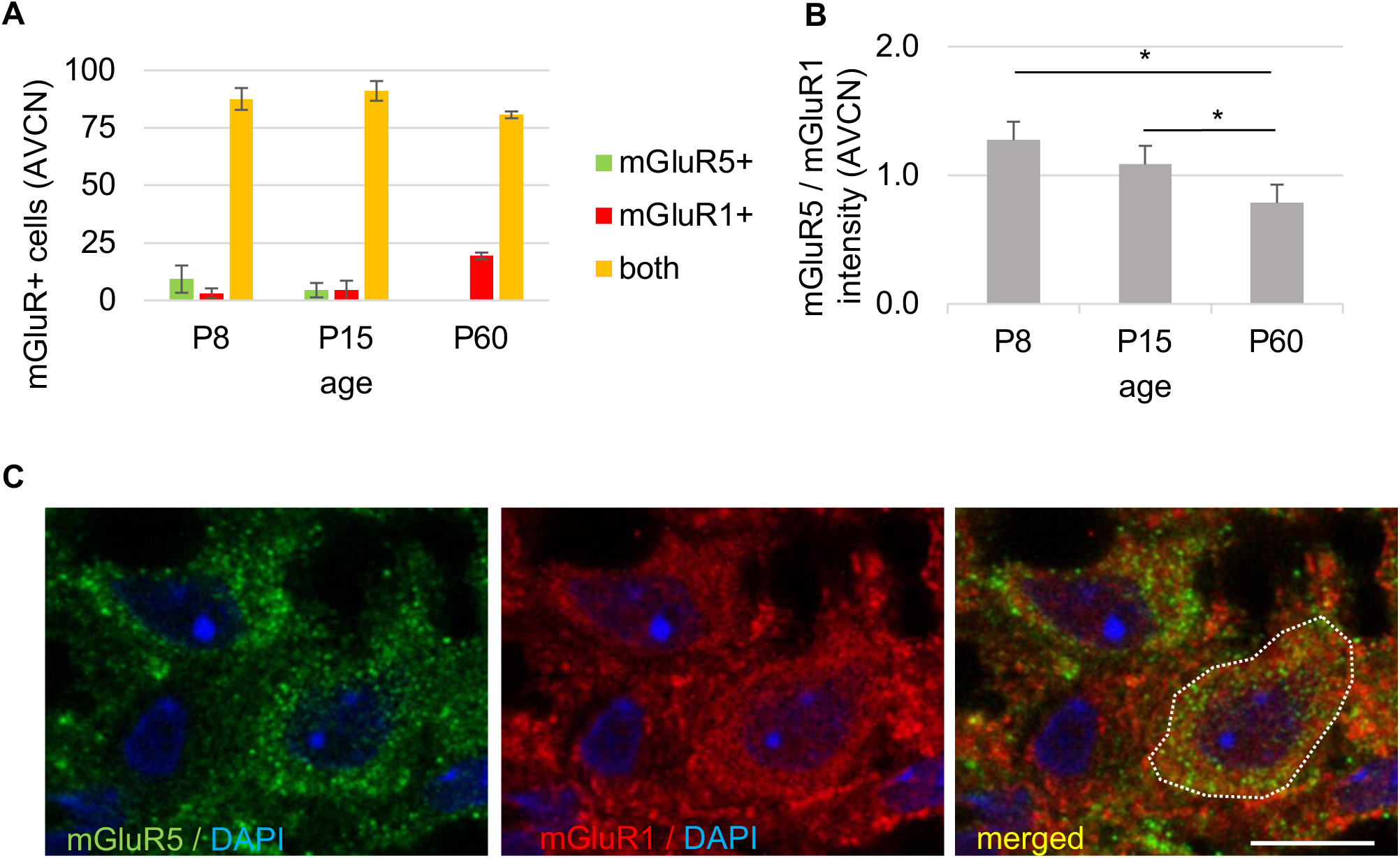
mGluR5 and mGluR1a are mostly co-expressed within the same cells in the AVCN (A) The majority ∼80+% of mGluR+ cells stain for both mGluR5 and mGluR1 at all ages (yellow bar). There is a small percentage of mGluR5+ only cells at P8 (green bar) and a small percentage that are mGluR1+ only at P60 (red bar). (B) The intensity of intracellular mGluR5 staining compared to mGluR1 within a cell declines with age. *p < 0.05 One-way ANOVA and posthoc Tukey test. (n = 3 mice). (C) Confocal imaging of the AVCN neurons at age P16 shows that mGluR5 (green) and mGluR1a (red) are mostly co-expressed within the same cell (example of a cell circumscribed with white dashed line). Scale bar = 10µm.

In-situ hybridization for *mGluR5* and *mGluR1* mRNA with RNAscope also demonstrates an age-dependent change from predominantly *mGluR5* to *mGluR1* expression with (**Fig. 7A**). Abundant co-expression of *mGluR5* and *mGluR1* is seen readily at P15 (**Fig. 7B**, dotted circles).

**Figure 7.**
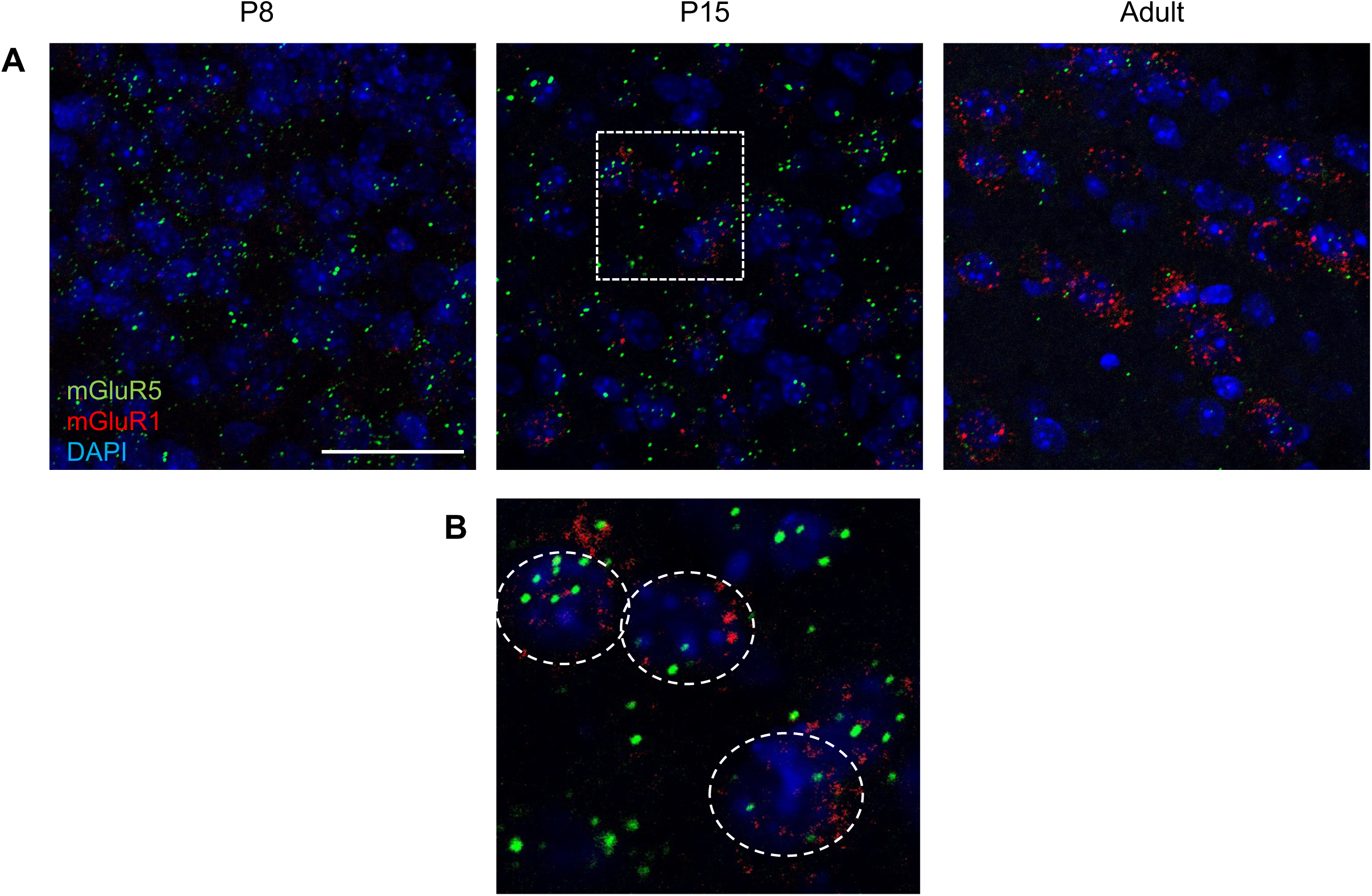
Co-expression and developmental shift from *mGluR5* to *mGluR1* mRNA expression in cochlear nucleus by RNAscope. (A) RNAscope in situ hybridization was performed on 14um cryosections through wildtype cochlear nucleus from ages P8, P15 and adult. Representative sections through ventral cochlear nucleus are shown for each age. There is also a developmental shift from primarily *mGluR5* (green) expression to *mGluR1* (red) expression with age, consistent with Western blotting and immunohistochemistry. (B) Enlarged boxed section from middle panel from (A) shows cells with co-expression of *mGluR5* and *mGluR1* (individual cells circumscribed). Scale bar = 10µm.

To identify which cochlear nucleus cell types express Group 1 mGluRs, we re-analyzed a single-cell transcriptomic dataset in which neuronal identity is correlated with electrophysiological properties using a Patch-seq platform (Jing et al., 2025). We independently performed UMAP and expression level analysis for each cell type cluster (using Seurat) and found that *mGluR5* and *mGluR1* are co-expressed in Bushy, Octopus, T-stellate, Fusiform and inhibitory neurons (**Fig. 8A-B)**. Violin plots show relative normalized expression across cell types (**Fig. 8C**). Consistent with the adult age of the animals in this dataset, the levels of *mGluR5* expression were substantially lower than *mGluR1* in most cell types. Fusiform neurons expressed the highest levels of *mGluR1* across all neuron types. Octopus cells exhibited a relatively higher expression of *mGluR5*. Collectively, the data indicate that Group1 *mGluRs* are co-expressed across most major neuron types in the cochlear nucleus.

**Figure 8.**
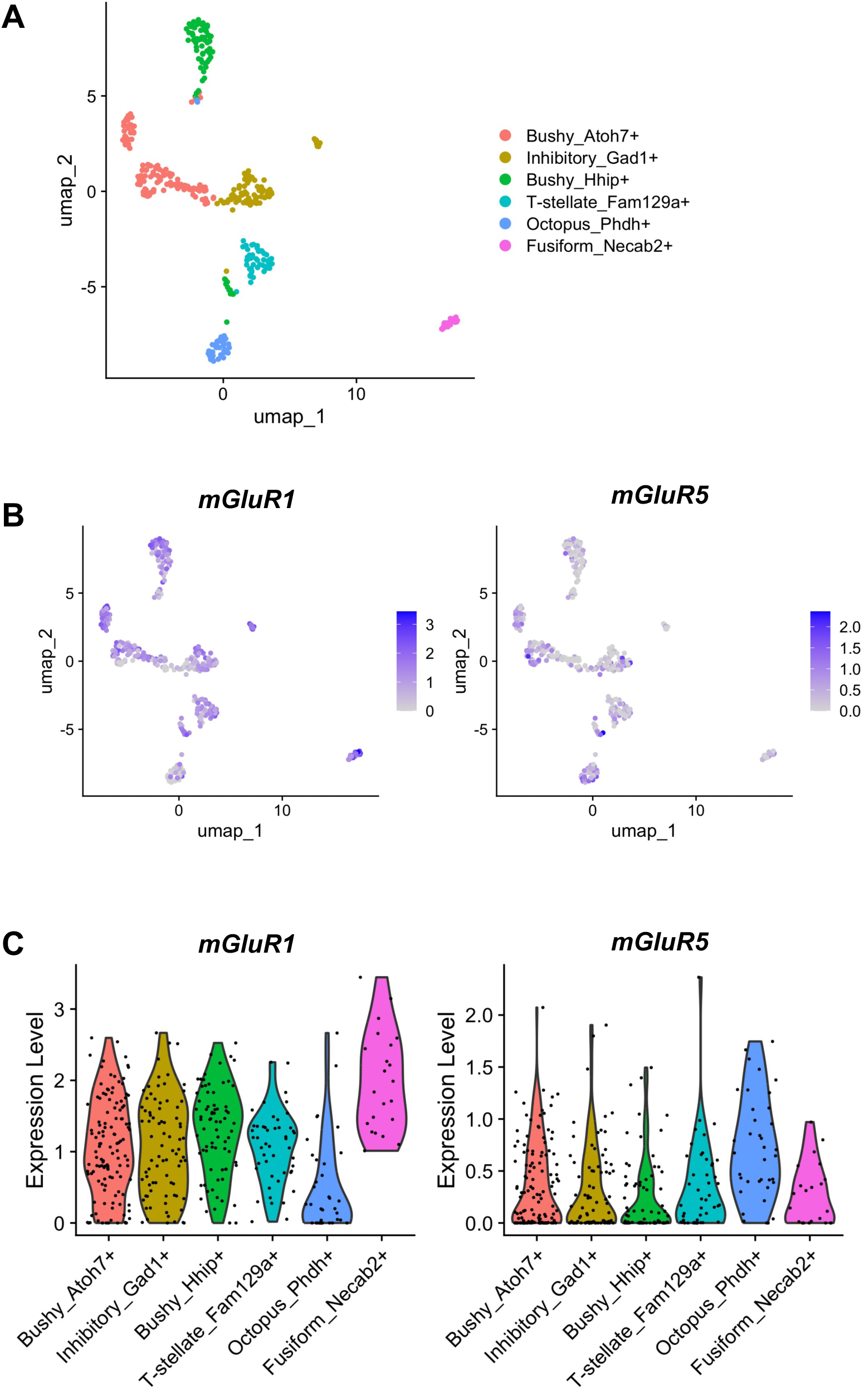
*mGluR1* and *mGluR5* co-expression across different neuron types in cochlear nucleus. Patch-seq data by Jing et al. 2025 was performed. (A) UMAP clustering of cell types shown with cell-specific markers. (B) There is overlapping distribution of *mGluR1* and *mGluR5* among multiple cell clusters. (C) Violin plot showing normalized relative expression of *mGluR1* and *mGluR5* across neuron types. Of note, *mGluR5* levels are relatively low due to the adult age of the tissue sample.

### Loss of cochlear activity did not affect the transition from *mGluR5* to *mGluR1* mRNA expression in both WT and *Fmr1* KO backgrounds

The mGluR5 to mGluR1 transition coincides with a time in development when mice transition from primarily spontaneous cochlear activity to sound evoked activity due to the opening of the ear canal and middle ear (Wang & Bergles, 2015). We sought to determine whether afferent (cochlear) activity is necessary for the transcriptional switch from *mGluR5* to *mGluR1* using three approaches **(Fig. 9A).** In the first experiment, we performed unilateral cochlea ablation in WT mice at P6 and assessed *mGluR1 and mGluR5* mRNA abundance in cochlear nucleus tissue at P16 using RT-qPCR. Cochlea ablation leads to an immediate cessation of cochlear innervation of cochlear nucleus. There was no difference in either *mGluR1* or *mGluR5* expression between the ablated and non-ablated sides (Paired *t*-test; n = 5 mice) (**Fig. 9B**). This indicates that sound-driven activity and cochlear innervation is not required for age-dependent changes in *mGluR5* or *mGluR1* expression.

**Figure 9.**
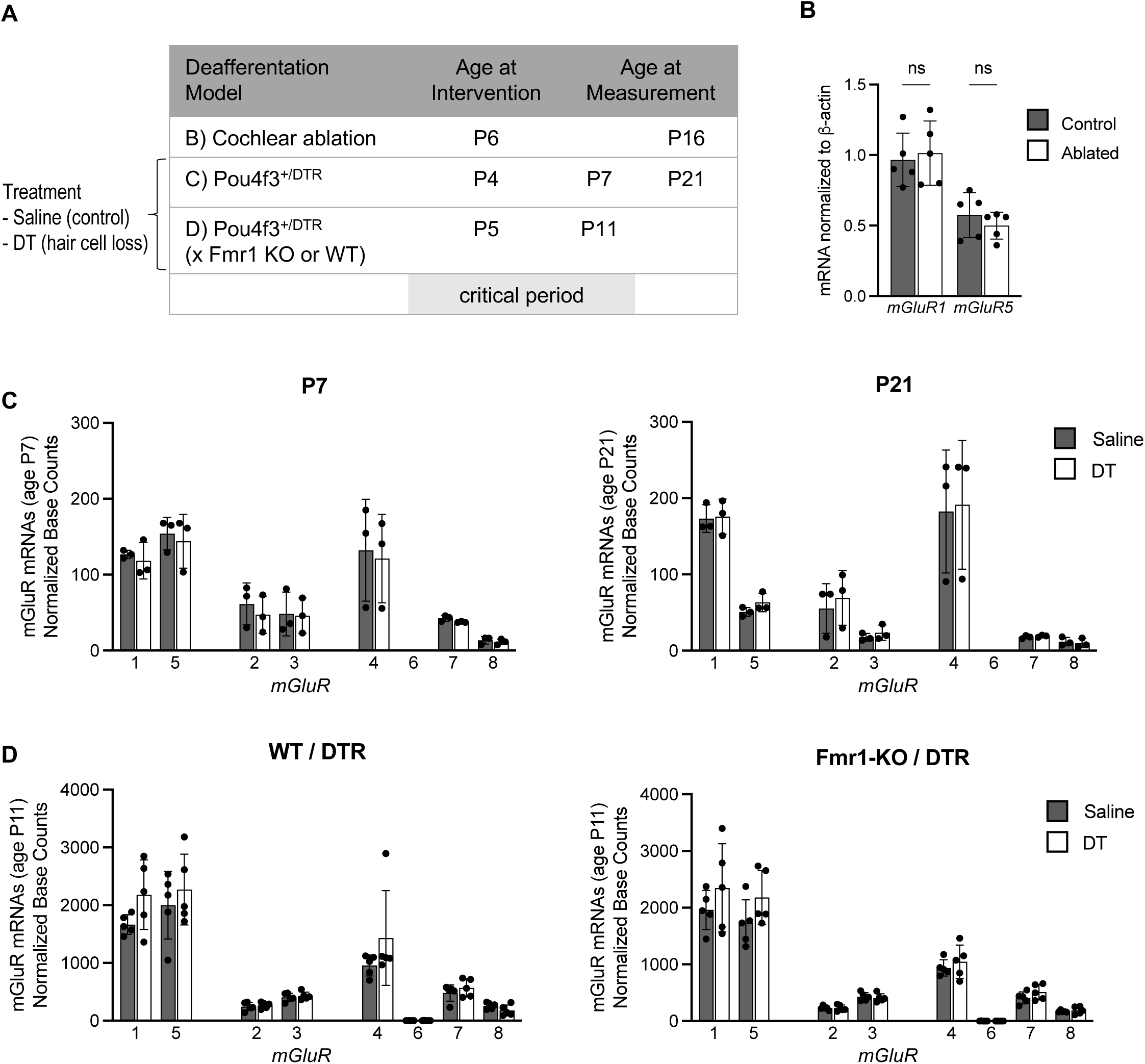
Loss of cochlear activity does not affect the transition from *mGluR5* to *mGluR1* mRNA expression over development. (A) Three models used to induce cochlear hair cell loss. (B) Unilateral cochlear ablation performed on WT mice at age P6 and cochlear nucleus collected at age P16. RT-qPCR for mGluR5 and mGluR1 shows no difference in ablated versus control side. (n = 5, paired t-test) (C) *Pou4f3*^+/DTR^ mice treated with saline or diphtheria toxin (DT) to induce cochlear hair cell loss at P2-5. Cochlear nuclei were harvested at P7 or P21. RNA-seq normalized base counts for mGluR1 and mGluR5 were no different between saline and DT but was statistically significantly different between ages. (n = 3, Holm-Šidák’s multiple comparisons test) (D) RNA-seq normalized base counts shown for *mGluR1* and *mGluR5* from P10 cochlear nuclei from *Pou4f3*^+/DTR^ mice crossed to *Fmr1* KO or WT background that were treated with saline (sal) or diphtheria toxin (DT) at P5-P10. (n = 5, Holm-Šidák’s multiple comparisons test).

In the second experiment, we utilized an inducible hair cell loss mouse model expressing diphtheria toxin receptor (DTR) under the *Pou4f3* promotor (*Pou4f3*^+/DTR^). This model has been previously characterized to exhibit substantial loss of hair cells and auditory brainstem responses by five days of diphtheria toxin (DT) administration (Tong et al., 2015). Mice were treated with either saline or DT at age P4, and cochlear nuclei were harvested at P7 and P21 for RNA-seq analysis (n = 3 biological replicates per age per treatment). Following either saline or DT injections, the levels of *mGluR1-8* transcript (normalized base count) changed with age, similar to the normal developmental pattern. There were no significant differences between saline-and DT-treated littermates at P7 (during the critical period) or at P21 (after the critical period) (**Fig. 9C**) (Holm-Šidák’s multiple comparison’s test), indicating that cochlear activity does not affect *mGluR1-8* expression in the cochlear nucleus.

Finally, in the last experiment, we asked whether FMRP has an influence in *mGluR1-8* expression under activity-deprived conditions. *Pou4f3*^+/DTR^ mice were crossed onto the *Fmr1-*KO background. Mice received a single DT administration at P5, and cochlear nuclei were collected at P11 for RNA-seq analysis (n = 5 biological replicates per genotype). There were no differences in *mGluR1-8* transcript levels between saline or DT treatment in WT or in *Fmr1* KO mice (**Fig. 9D**) (Holm-Šidák’s multiple comparison’s test). These findings indicate that FMRP does not influence the activity-independent nature of the *mGluR1-8* gene expression or the developmental transition in *mGluR5-mGluR1* transcripts.

### Gene ontology reveals differentially expressed genes in *Fmr1* KO are enriched for glutamate receptor signaling

While we did not find significant differences in the expression levels of metabotropic glutamate receptors between *Fmr1* KO and WT cochlear nucleus, gene ontology analysis of differentially expressed genes (DEGs) at age P8 support the idea that the glutamatergic pathways may be dysregulated. Among differentially expressed genes (p_adj_ < 0.05, log_2_ fold change > 0.1) with sufficient expression level (base mean count >100) were 36 upregulated genes and 49 down regulated genes in *Fmr1* KO. There were no significantly enriched pathways with upregulated genes due to the low number of genes. However, among downregulated genes, Syngo analysis identified an enrichment for postsynapse cellular component (**Fig. 10A**). DAVID analysis yielded similar results with the top enrichment for the glutamatergic synapse and postsynapse (**Fig. 10B**). Downregulated DEGs within this pathway are listed **(Fig. 10C)**. Notably, one downregulated gene is *Prkar1b*, a subunit of PKA kinase known to phosphorylate mGluR5 but not mGluR1 (Uematsu et al., 2015).

**Figure 10.**
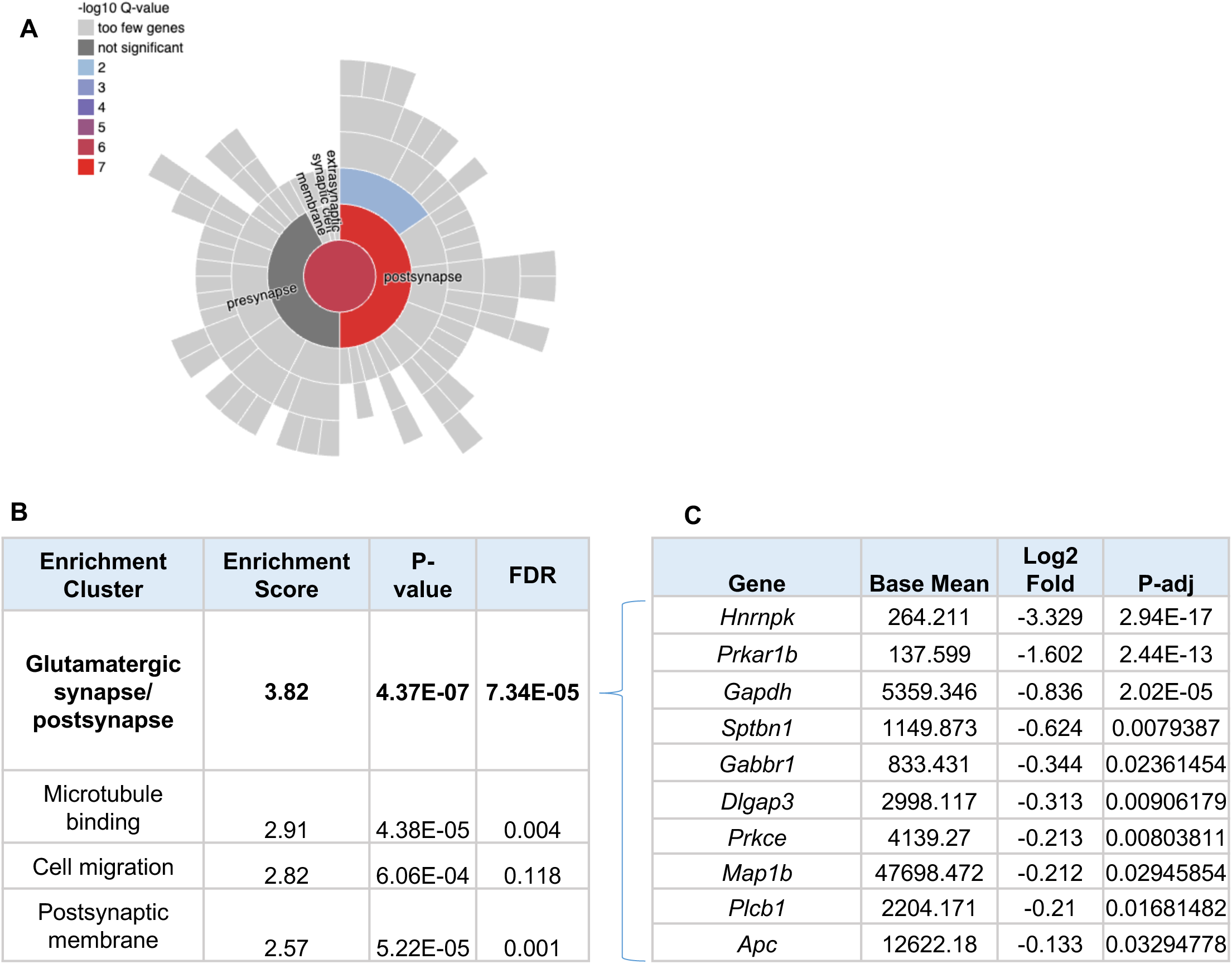
Gene ontology analysis of downregulated cochlear nucleus genes in *Fmr1* KO at P8. (A) Syngo analysis of 49 downregulated genes show clustering of genes localized to the postsynapse. (B) DAVID analysis reveals clustering of 10 genes involved in glutamatergic postsynapse. (C) List of genes clustered in the postsynapse group from (B) and their relative expression in *Fmr1* KO compared to WT cochlear nucleus.

## Discussion

This study conducted multiple mRNA and protein analyses on tissue samples specifically collected from the cochlear nucleus in mice across ages and *Fmr1* genotypes. The results, as summarized in **Fig. 11**, lead to three novel findings. First, we identified a developmental transition in the cochlear nucleus, marked by mGluR5 expression level starting high and decreasing with age, and by mGluR1 expression starting low and progressively increasing. The transition takes place in the second postnatal week in mice. Second, there appears to be co-expression of mGluR5 and mGluR1 among various neurons in the cochlear nucleus. Third, the transition from mGluR5 to mGluR1 gene expression is an intrinsic, developmental event that is independent of afferent cochlear activity under both WT and *Fmr1* KO conditions.

**Figure 11.**
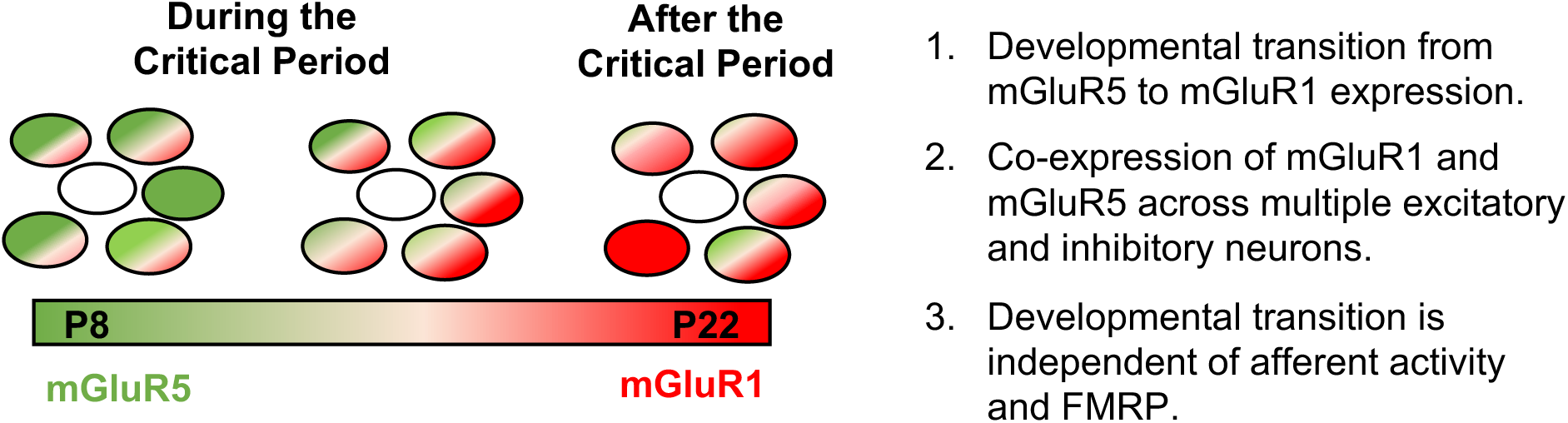
Schematic of cochlear nucleus neurons expressing mGluR5 and mGluR1 with age.

### Developmental transition from mGluR5 to mGluR1 expression

This is the first report of a differential developmental expression of group I *mGluR* expression in the cochlear nucleus. It is known that *mGluR1* is the predominant *mGluR* expressed in the adult cochlear nucleus (Petralia et al., 2000). We confirmed this observation at both the transcript and protein levels. More importantly, and not previously known, we identified that *mGluR5* is highly expressed in cochlear nucleus in early postnatal development. The most dramatic changes in *mGluR1* or *mGluR5* levels occurred between P8 and P22. The mGluR5/mGluR1 ratio declines within these cells with age.

One functional implication of this transition may be related to the closure of auditory critical periods. The most dramatic changes in *mGluR1* or *mGluR5* levels occurred between P8 and P22, a period during which a well-studied critical period in the VCN closes. The auditory critical period in mouse cochlear nucleus has been defined by the survival response of AVCN neurons to afferent deprivation using cochlear ablation methods (Mostafapour et al., 2000). It is known that deafferentation during the critical period causes massive cell death in the AVCN between P5 and P10, whereas the same procedure performed after the critical period (>P14) shows minimal cell death. The change in mGluR5/mGluR1 levels occurs during the closure of the critical period when AVCN neurons become less susceptible to apoptosis with ipsilateral cochlear deafferentation. (Mostafapour et al., 2000) Since pro-survival pathways such as mTOR and ERK/MAPK can be activated by group I mGluRs, it would be interesting to determine in future experiments if the mGluR1 and mGluR5 confer different survivability of AVCN neurons in response to loss of afferent activity.

The closure of the critical period in AVCN occurs by postnatal day 14. What triggers this change in neuroplasticity is not well understood. During the critical period, afferent activity is provided by spontaneous cochlear activity (Kros et al., 1998). Hearing onset occurs around the time of critical period closure after which afferent activity is provided by sound evoked responses. (Anthwal & Thompson, 2016) In our DTR and cochlear ablation hearing loss models, we show that the loss of cochlear afferent activity, which affect both spontaneous activity and evoked activity, are not necessary for the transition of *mGluR* gene expression. Thus, the change in *mGluR* mRNA expression is intrinsic to cochlear nucleus neuron maturity.

A potential link between mGluR expression dynamics and sensory critical period properties is supported by studies from other systems. Changes in mGluR5 expression during development has been described in other sensory systems, such as the cat visual cortex where there is significant decline during peak critical period (Reid et al., 1995). There are also changes in the laminar distribution of expression of mGluR5 which is delayed with visual deprivation.

### What is the functional relevance of mGluR5/mGluR1 dynamics

Both mGluR1 and mGluR5 have been considered functionally redundant due to their sequence similarity, shared coupling to Gα_q/11_ and activation of PLC pathways, and response to the agonist 3,5-dihydroxyphenylglycine (DHPG). In support of this notion, behavioral abnormalities in the mGluR1 KO mouse can be rescued by mGluR5 expression in cerebellar Purkinje cells (Harbers et al., 2022). On the other hand, several studies highlight functional divergence between mGluR1 and mGluR5 (Valenti et al., 2002). Activation of protein kinase C differentially regulates mGluR1α and mGluR5α, selectively desensitizing intracellular Ca² mobilization in mGluR1α-expressing HEK-293 cells while sustaining oscillatory Ca² mobilization in mGluR5α-expressing cells, highlighting distinct coupling mechanisms during prolonged mGluR stimulation (Kawabata et al., 1998). Both mGluR receptors have splice variants, and while both mGluR5 splice variants can interact with Homer (a scaffolding protein needed for IP3 signaling), the mGluR1α splice variant lacks the C-terminal tail which interacts with Homer. (Brakeman et al., 1997; Tu et al., 1998) In CA1 hippocampal neurons that endogenously express both receptors, DHPG-induced calcium responses are mediated by mGluR1, whereas DHPG-induced after-hyperpolarization and NMDA potentiation are mediated by mGluR5 (Mannaioni et al., 2001). Additionally, although activation of either mGluR1 or mGluR5 alone can induce long-term depression (LTD) in hippocampal neurons, only mGluR1 reduces surface AMPA receptor levels (Volk et al., 2006). In the basal ganglia, where both receptors are expressed, mGluR1 mediates DPHG-induced depolarization, while mGluR5 antagonism blocks the desensitization of mGluR1-mediated depolarization (Poisik et al., 2003). In the subthalamic nucleus, both receptors are abundantly expressed postsynaptically but only mGluR5 mediates DHPG-induced activation (Awad et al., 2000). PKA kinase can phosphorylate mGluR5 but not mGluR1 (Uematsu et al., 2015). Collectively, these studies suggest that the two Group 1 mGluRs have overlapping major functions but distinct roles in fine-tuned regulation. It is possible that mGluR1 and mGluR5 may differentially modulate different aspects of neuron and synapse properties, and the developmental shift may due to functional needs of the developing cochlear nucleus.

Additionally, cellular expression and subcellular localization of mGluR receptors should be considered with regards to the functional implication of the mGluR5-mGluR1 transition. The cochlear nucleus is comprised of several different cell types which were recently characterized by patch-seq with discovery of molecular markers (Jing et al., 2025). In our analysis of this publicly available data, we found mGluR5 and mGluR1 are broadly expressed in multiple cell types. However, further analysis on the cellular expression of group I mGluR over development is needed. At the subcellular level, mGluR1 is mainly postsynaptic both during development and in adulthood. (Petralia et al., 1996) No published studies on mGluR5 localization in the cochlear nucleus are currently available. In another auditory brainstem nucleus MNTB, mGluR5 is localized both pre-and post-synaptically (Peng et al., 2020; Wang et al., 2023). Further investigations on the localization patterns of mGluR1 and mGluR5 in the cochlear nucleus are needed to address how the transition from predominantly mGluR5 to mGluR1 may affect physiological changes in cochlear nucleus neurons during development.

### mGluR5 protein level in the absence of FMRP

Abnormal mGluR5 signal has long been associated with *Fmr1* KO-induced neural alterations in mature systems, primarily attributed to the loss of FMRP regulation over signals downstream of mGluR5 activation (Bear et al., 2004), rather than direct alterations in mGluRs. While many abnormalities of *Fmr1* KO can be corrected by inhibiting mGluR5 function, there is no consistent evidence that mGluR5 protein levels are changed. This notion is supported by the failure to identify a change in the overall level of mGluR5 by western blotting across several forebrain areas and cerebellum throughout the development of *Fmr1* KO mice (Giuffrida et al., 2005), although one study reported a marginal but significant mGluR5 increase in the prefrontal cortex of individuals with FXS (Lohith et al., 2013). It is important to note that changes in mGluR5 protein level may not necessarily correlate with receptor availability. Elevations in mGluR5 availability as measured by [^3^H]-labeled 3-methoxy-5-pyridin-2-ylethynylpyridine (MPEPy) radioligand binding were not statistically significant in FXS (Lohith et al., 2013). Intriguingly, a more recent study using positron emission tomography (PET) imaging reported lower 3-[18F]fluoro-5-(2-pyridinylethynyl)benzonitrile ([18F] (FPEB) binding potential, reflecting reduced mGluR5 availability in adult individuals with FXS (Mody et al., 2021; Brašić et al., 2022). The binding potential similarly reduces in adult *Fmr1* KO mice, and interestingly, prior to this reduction, the binding potential increases in young males but not females (Afshar et al., 2022). In this study, did not find a statistically significant difference in mGluR5 protein level in *Fmr1* KO compared to WT among sexes and strains. While transcriptome differences were small between *Fmr1* KO and WT cochlear nuclei, it is interesting that the downregulated genes were enriched for glutamatergic postsynapse. This may be a result of feedback from dysregulated mGluR5 signaling, which warrant further investigation.

In summary, we discover a developmental transition from predominantly mGluR5 to mGluR1 expression that occurs during the closure of the critical period of cochlear nucleus development. This transition appears to be independent of afferent cochlear activity. Although we did not find statistically significant differences in mGluR expression between *Fmr1* KO and WT, there were transcriptomic differences among genes related to the glutamatergic post synapse which may reflect abnormalities in the mGluR signaling pathways.

## Acknowledgements

We thank Dr. Lynne Maquat, Dr. Kimberly Huber and their lab members for helpful discussions; Dr. Laurent Gatron for assistance with the RNAscope; Ling Tong for assistance with DT injection and tissue dissection; and Dr. Deborah Nickerson, Dr. JW MacDonald, Dr. John Ashton and colleagues for assistance with the RNA sequencing.

## Key Points

Group 1 mGluRs are expressed in multiple neuron types in the cochlear nucleus. There is a transition from predominantly mGluR5 to mGluR1 expression during the auditory critical period closure which appears to be independent of cochlear activity or absence of FMRP.

## Author Contributions

P.K. and J.L. performed immunostaining; J.W. and J.K. performed western blotting; M.C. performed the RT-qPCR; X.Y. and R.G. performed the experiments with Pou4f3+/DTR; E.R., Y.W. and H.S. performed the study design, experiments, analysis, and manuscript preparation.

## Data Availability Statement

The RNA-seq datasets (raw and processed) for this study have been uploaded onto the NCBI GEO repository (https://www.ncbi.nlm.nih.gov/geo/) and will be made publicly available at the time of publication. See Methods for accession numbers.

## References

Afshar, S., Lule, S., Yuan, G., Qu, X., Pan, C., Whalen, M., Brownell, A. L., & Mody, M. (2022). Longitudinal PET studies of mGluR5 in FXS using an FMR1 knockout mouse model. Transl Neurosci, 13(1), 80–92. 10.1515/tnsci-2022-0217

Anthwal, N., & Thompson, H. (2016). The development of the mammalian outer and middle ear.J Anat, 228(2), 217–232. 10.1111/joa.12344

Awad, H., Hubert, G. W., Smith, Y., Levey, A. I., & Conn, P. J. (2000). Activation of metabotropic glutamate receptor 5 has direct excitatory effects and potentiates NMDA receptor currents in neurons of the subthalamic nucleus. J Neurosci, 20(21), 7871–7879. 10.1523/JNEUROSCI.20-21-07871.2000

Bilak, S. R., & Morest, D. K. (1998). Differential expression of the metabotropic glutamate receptor mGluR1alpha by neurons and axons in the cochlear nucleus: in situ hybridization and immunohistochemistry. Synapse, 28(4), 251–270. 10.1002/(SICI)1098-2396(199804)28:4<251::AID-SYN1>3.0.CO;2-8

Brakeman, P. R., Lanahan, A. A., O’Brien, R., Roche, K., Barnes, C. A., Huganir, R. L., & Worley, P. F. (1997). Homer: a protein that selectively binds metabotropic glutamate receptors. Nature, 386(6622), 284–288. 10.1038/386284a0

Chen, L., & Toth, M. (2001). Fragile X mice develop sensory hyperreactivity to auditory stimuli. Neuroscience, 103(4), 1043–1050. 10.1016/s0306-4522(01)00036-7

Consortium, T. D.-B. F. X. (1994). Fmr1 knockout mice: a model to study fragile X mental retardation. The Dutch-Belgian Fragile X Consortium. Cell, 78(1), 23–33. https://www.ncbi.nlm.nih.gov/pubmed/8033209

Dahl, V., Helmbrecht, H., Rios Sigler, A., Hildahl, K., Sullivan, H., Janakiraman, S., Jasti, S., & Nance, E. (2022). Characterization of a mGluR5 Knockout Rat Model with Hallmarks of Fragile X Syndrome. Life (Basel), 12(9). 10.3390/life12091308

Dolen, G., & Bear, M. F. (2008). Role for metabotropic glutamate receptor 5 (mGluR5) in the pathogenesis of fragile X syndrome. J Physiol, 586(6), 1503–1508. 10.1113/jphysiol.2008.150722

Dolen, G., Osterweil, E., Rao, B. S., Smith, G. B., Auerbach, B. D., Chattarji, S., & Bear, M. F. (2007). Correction of fragile X syndrome in mice. Neuron, 56(6), 955–962. 10.1016/j.neuron.2007.12.001

Frankland, P. W., Wang, Y., Rosner, B., Shimizu, T., Balleine, B. W., Dykens, E. M., Ornitz, E. M., & Silva, A. J. (2004). Sensorimotor gating abnormalities in young males with fragile X syndrome and Fmr1-knockout mice. Mol Psychiatry, 9(4), 417–425. 10.1038/sj.mp.4001432

Giuffrida, R., Musumeci, S., D’Antoni, S., Bonaccorso, C. M., Giuffrida-Stella, A. M., Oostra, B. A., & Catania, M. V. (2005). A reduced number of metabotropic glutamate subtype 5 receptors are associated with constitutive homer proteins in a mouse model of fragile X syndrome. J Neurosci, 25(39), 8908–8916. 10.1523/JNEUROSCI.0932-05.2005

Gonzalez, D., Tomasek, M., Hays, S., Sridhar, V., Ammanuel, S., Chang, C. W., Pawlowski, K., Huber, K. M., & Gibson, J. R. (2019). Audiogenic Seizures in the Fmr1 Knock-Out Mouse Are Induced by Fmr1 Deletion in Subcortical, VGlut2-Expressing Excitatory Neurons and Require Deletion in the Inferior Colliculus. J Neurosci, 39(49), 9852–9863. 10.1523/JNEUROSCI.0886-19.2019

Harbers, M., Nakao, H., Watanabe, T., Matsuyama, K., Tohyama, S., Nakao, K., Kishimoto, Y., Kano, M., & Aiba, A. (2022). mGluR5 Is Substitutable for mGluR1 in Cerebellar Purkinje Cells for Motor Coordination, Developmental Synapse Elimination, and Motor Learning. Cells, 11(13). 10.3390/cells11132004

Holley, A. J., Shedd, A., Boggs, A., Lovelace, J., Erickson, C., Gross, C., Jankovic, M., Razak, K., Huber, K., & Gibson, J. R. (2022). A sound-driven cortical phase-locking change in the Fmr1 KO mouse requires Fmr1 deletion in a subpopulation of brainstem neurons. Neurobiol Dis, 170, 105767. 10.1016/j.nbd.2022.105767

Huang da, W., Sherman, B. T., & Lempicki, R. A. (2009). Systematic and integrative analysis of large gene lists using DAVID bioinformatics resources. Nat Protoc, 4(1), 44–57. 10.1038/nprot.2008.211

Huber, K. M., Gallagher, S. M., Warren, S. T., & Bear, M. F. (2002). Altered synaptic plasticity in a mouse model of fragile X mental retardation. Proc Natl Acad Sci U S A, 99(11), 7746–7750. 10.1073/pnas.122205699

Jing, J., Hu, M., Ngodup, T., Ma, Q., Lau, S. N., Ljungberg, M. C., McGinley, M. J., Trussell, L. O., & Jiang, X. (2025). Molecular logic for cellular specializations that initiate the auditory parallel processing pathways. Nat Commun, 16(1), 489. 10.1038/s41467-024-55257-z

Kawabata, S., Kohara, A., Tsutsumi, R., Itahana, H., Hayashibe, S., Yamaguchi, T., & Okada, M. (1998). Diversity of calcium signaling by metabotropic glutamate receptors. J Biol Chem, 273(28), 17381–17385. 10.1074/jbc.273.28.17381

Kros, C. J., Ruppersberg, J. P., & Rusch, A. (1998). Expression of a potassium current in inner hair cells during development of hearing in mice. Nature, 394(6690), 281–284. 10.1038/28401

Litovchick, L. (2020). Immunoblotting. Cold Spring Harb Protoc, 2020(6), 098392. 10.1101/pdb.top098392

Lohith, T. G., Osterweil, E. K., Fujita, M., Jenko, K. J., Bear, M. F., & Innis, R. B. (2013). Is metabotropic glutamate receptor 5 upregulated in prefrontal cortex in fragile X syndrome? Mol Autism, 4(1), 15. 10.1186/2040-2392-4-15

Lu, Y. (2014). Metabotropic glutamate receptors in auditory processing. Neuroscience, 274, 429–445. 10.1016/j.neuroscience.2014.05.057

Mannaioni, G., Marino, M. J., Valenti, O., Traynelis, S. F., & Conn, P. J. (2001). Metabotropic glutamate receptors 1 and 5 differentially regulate CA1 pyramidal cell function. J Neurosci, 21(16), 5925–5934. 10.1523/JNEUROSCI.21-16-05925.2001

McCullagh, E. A., Rotschafer, S. E., Auerbach, B. D., Klug, A., Kaczmarek, L. K., Cramer, K. S., Kulesza, R. J., Jr., Razak, K. A., Lovelace, J. W., Lu, Y., Koch, U., & Wang, Y. (2020). Mechanisms underlying auditory processing deficits in Fragile X syndrome. FASEB J, 34(3), 3501–3518. 10.1096/fj.201902435R

Michalon, A., Sidorov, M., Ballard, T. M., Ozmen, L., Spooren, W., Wettstein, J. G., Jaeschke, G., Bear, M. F., & Lindemann, L. (2012). Chronic pharmacological mGlu5 inhibition corrects fragile X in adult mice. Neuron, 74(1), 49–56. 10.1016/j.neuron.2012.03.009

Mody, M., Petibon, Y., Han, P., Kuruppu, D., Ma, C., Yokell, D., Neelamegam, R., Normandin, M. D., Fakhri, G. E., & Brownell, A. L. (2021). In vivo imaging of mGlu5 receptor expression in humans with Fragile X Syndrome towards development of a potential biomarker. Sci Rep, 11(1), 15897. 10.1038/s41598-021-94967-y

Mostafapour, S. P., Cochran, S. L., Del Puerto, N. M., & Rubel, E. W. (2000). Patterns of cell death in mouse anteroventral cochlear nucleus neurons after unilateral cochlea removal. J Comp Neurol, 426(4), 561–571. 10.1002/1096-9861(20001030)426:4<561::aid-cne5>3.0.co;2-g

Nielsen, D. M., Derber, W. J., McClellan, D. A., & Crnic, L. S. (2002). Alterations in the auditory startle response in Fmr1 targeted mutant mouse models of fragile X syndrome. Brain Res, 927(1), 8–17. 10.1016/s0006-8993(01)03309-1

Niswender, C. M., & Conn, P. J. (2010). Metabotropic glutamate receptors: physiology, pharmacology, and disease. Annu Rev Pharmacol Toxicol, 50, 295–322. 10.1146/annurev.pharmtox.011008.145533

Petralia, R. S., Rubio, M. E., Wang, Y. X., & Wenthold, R. J. (2000). Differential distribution of glutamate receptors in the cochlear nuclei. Hear Res, 147(1-2), 59–69. 10.1016/s0378-5955(00)00120-9

Petralia, R. S., Wang, Y. X., Zhao, H. M., & Wenthold, R. J. (1996). Ionotropic and metabotropic glutamate receptors show unique postsynaptic, presynaptic, and glial localizations in the dorsal cochlear nucleus. J Comp Neurol, 372(3), 356–383. 10.1002/(SICI)1096-9861(19960826)372:3<356::AID-CNE3>3.0.CO;2-1

Poisik, O. V., Mannaioni, G., Traynelis, S., Smith, Y., & Conn, P. J. (2003). Distinct functional roles of the metabotropic glutamate receptors 1 and 5 in the rat globus pallidus. J Neurosci, 23(1), 122–130. 10.1523/JNEUROSCI.23-01-00122.2003

Reid, S. N., Romano, C., Hughes, T., & Daw, N. W. (1995). Immunohistochemical study of two phosphoinositide-linked metabotropic glutamate receptors (mGluR1 alpha and mGluR5) in the cat visual cortex before, during, and after the peak of the critical period for eye-specific connections. J Comp Neurol, 355(3), 470–477. 10.1002/cne.903550311

Rotschafer, S. E., & Razak, K. A. (2014). Auditory processing in fragile x syndrome. Front Cell Neurosci, 8, 19. 10.3389/fncel.2014.00019

Santoro, M. R., Bray, S. M., & Warren, S. T. (2012). Molecular mechanisms of fragile X syndrome: a twenty-year perspective. Annu Rev Pathol, 7, 219–245. 10.1146/annurev-pathol-011811-132457

Schneider, C. A., Rasband, W. S., & Eliceiri, K. W. (2012). NIH Image to ImageJ: 25 years of image analysis. Nat Methods, 9(7), 671–675. 10.1038/nmeth.2089

Tang, Z. Q., & Lu, Y. (2018). Anatomy and Physiology of Metabotropic Glutamate Receptors in Mammalian and Avian Auditory System. HSOA Trends Anat Physiol, 1. 10.24966/TAP-7752/100001

Thomazeau, A., Bosch, M., Essayan-Perez, S., Barnes, S. A., De Jesus-Cortes, H., & Bear, M. F. (2021). Dissociation of functional and structural plasticity of dendritic spines during NMDAR and mGluR-dependent long-term synaptic depression in wild-type and fragile X model mice. Mol Psychiatry, 26(9), 4652–4669. 10.1038/s41380-020-0821-6

Tong, L., Strong, M. K., Kaur, T., Juiz, J. M., Oesterle, E. C., Hume, C., Warchol, M. E., Palmiter, R. D., & Rubel, E. W. (2015). Selective deletion of cochlear hair cells causes rapid age-dependent changes in spiral ganglion and cochlear nucleus neurons. J Neurosci, 35(20), 7878–7891. 10.1523/JNEUROSCI.2179-14.2015

Tu, J. C., Xiao, B., Yuan, J. P., Lanahan, A. A., Leoffert, K., Li, M., Linden, D. J., & Worley, P. F. (1998). Homer binds a novel proline-rich motif and links group 1 metabotropic glutamate receptors with IP3 receptors. Neuron, 21(4), 717–726. 10.1016/s0896-6273(00)80589-9

Uematsu, K., Heiman, M., Zelenina, M., Padovan, J., Chait, B. T., Aperia, A., Nishi, A., & Greengard, P. (2015). Protein kinase A directly phosphorylates metabotropic glutamate receptor 5 to modulate its function. J Neurochem, 132(6), 677–686. 10.1111/jnc.13038

Valenti, O., Conn, P. J., & Marino, M. J. (2002). Distinct physiological roles of the Gq-coupled metabotropic glutamate receptors Co-expressed in the same neuronal populations. J Cell Physiol, 191(2), 125–137. 10.1002/jcp.10081

Volk, L. J., Daly, C. A., & Huber, K. M. (2006). Differential roles for group 1 mGluR subtypes in induction and expression of chemically induced hippocampal long-term depression. J Neurophysiol, 95(4), 2427–2438. 10.1152/jn.00383.2005

Wang, H. C., & Bergles, D. E. (2015). Spontaneous activity in the developing auditory system. Cell Tissue Res, 361(1), 65–75. 10.1007/s00441-014-2007-5

Yan, Q. J., Rammal, M., Tranfaglia, M., & Bauchwitz, R. P. (2005). Suppression of two major Fragile X Syndrome mouse model phenotypes by the mGluR5 antagonist MPEP. Neuropharmacology, 49(7), 1053–1066. 10.1016/j.neuropharm.2005.06.004

Yu, X., & Wang, Y. (2023). Peripheral Fragile X messenger ribonucleoprotein is required for the timely closure of a critical period for neuronal susceptibility in the ventral cochlear nucleus. Front Cell Neurosci, 17, 1186630. 10.3389/fncel.2023.1186630

Yun, S. W., Platholi, J., Flaherty, M. S., Fu, W., Kottmann, A. H., & Toth, M. (2006). Fmrp is required for the establishment of the startle response during the critical period of auditory development. Brain Res, 1110(1), 159–165. 10.1016/j.brainres.2006.06.086

